# The global spread and invasion capacities of alien ants

**DOI:** 10.1101/2021.11.19.469299

**Authors:** Mark K. L. Wong, Evan P. Economo, Benoit Guénard

## Abstract

The ecological and economic impacts of biological invasions are usually highly conspicuous, but these are the outcome of a global, multistage process that is obscured from view. For most taxa, we lack a large-scale picture of the movements of alien species, the biases and barriers that promote or inhibit their spread at each stage, and blind spots in our ability to detect species during their spread. For instance, countries rely heavily on customs interceptions to prevent new species introductions, but their efficacy for detecting invaders remains unclear. To address these gaps, we synthesize and provide data at unprecedented geographic resolution on the global diversity and distribution of alien ants, a pervasive group strongly impacting humans and ecosystems. From >145,000 records spanning 602 regions, we identify 522 ant species exhibiting human-mediated spread, doubling recent estimates of their diversity. We show that movements of alien ant species across regions globally are non-random and, moreover, that these flows differ by the extents to which species invade—ranging from arrival to indoor establishment, naturalization, and harmful status. Importantly, we find that almost two-thirds of the 309 species that naturalize globally—most of which are ground- and litter-dwelling—are absent from customs interceptions, which record disproportionately high numbers of arboreal species. Our results illustrate the vast, yet uneven extent of ant invasions globally, and suggest that most alien species bypass biosecurity controls while spreading successfully worldwide. This raises doubts on the efficacy of current customs interceptions procedures and highlights a need for radically new approaches.

**Significance statement:** Biological invasions impact humans and ecosystems tremendously. Invasions are difficult to address because little is known about large-scale patterns of spread, species’ capacities to invade ecosystems, and the efficacy of existing biosecurity measures. This paper tackles these issues for alien ants, one of the most damaging groups of invasive animals. An unprecedented dataset reveals that 522 ant species exhibit human-mediated spread, 270 more than previously known. Species are grouped under four levels of invasion capacity corresponding to different invasion barriers. Geographic sources and sinks for the groups differ globally. Two-thirds of species with naturalization capacity have never been recorded at customs interceptions, which fail to detect many litter-dwelling species. Novel detection and control strategies for inconspicuous alien organisms are needed.

## INTRODUCTION

Incessant movements of humans and goods are driving an unprecedented exchange of organisms worldwide, and the ensuing biological invasions cost the global economy US$26.8 billion annually (Elton, 1958; Seebens et al., 2017; Diagne et al., 2021). The spread and impacts of alien species introduced deliberately for economic or cultural purposes, such as various ornamental plants and mammalian predators, are relatively well studied (Doherty et al., 2016; Van Kleunen et al., 2020). However, barring a few exceptions (Kraemer et al., 2015), there is little information about the global spread of the great majority of alien species that stowaway on transport and that are introduced unintentionally (Turbelin et al., 2017). Furthermore, the extent to which such species, especially numerous invertebrates, plants, and fungi, are detected in customs interceptions—which provide vital gatekeeping functions for the biosecurity of most countries—remains unclear.

As not all alien species pose equal threats, effective management of invasions requires accurate prediction of species’ capacities to naturalize and cause harm. From an ecological perspective, this “invasion capacity” of a species is determined by the extent to which its individuals or propagules can overcome geographical barriers to their transport, and—upon subsequent arrival—overcome intrinsic demographic- and extrinsic environmental barriers to their establishment (survival and reproduction) and spread of viable populations (Blackburn et al., 2011). Although geographic patterns of species exchange have been documented at the global scale for several groups (Van Kleunen et al., 2015; Dawson et al., 2017), differences among species’ invasion capacities are seldom addressed. Most studies focus only on “naturalized” or “invasive” species—subsets of alien species that can overcome all barriers discussed above (Van Kleunen et al., 2015; Dawson et al., 2017). Other studies have treated records of alien species that are unique to customs interceptions or indoor settings similarly to records of those with established populations in the wild (Miravete et al., 2014; Bertelsmeier et al., 2017, Bertelsmeier & Ollier, 2021). Such approaches overlook fundamental differences in invasion outcomes and may mischaracterize global patterns of invasions. Moreover, they miss key opportunities to understand the mechanisms responsible for species’ differing invasion capacities and to develop targeted biosecurity measures in response.

Ants are archetypal stowaway organisms (Weber, 1939) and excellent models for understanding the global spread of unintentionally introduced species. Many alien ants naturalize, and multiple cause severe ecological, economic, and social damages globally (Holway et al., 2002; Diagne et al., 2021). Records of ant species’ occurrences across regions worldwide have also been consolidated recently (Guénard et al., 2017). Nonetheless, there have been no attempts to investigate the global spread of alien ant species while explicitly accounting for species’ invasion capacities. For instance, in previous studies summarizing the diversity and distribution of alien ants (McGlynn, 1999; Suarez et al., 2010; Miravete et al., 2014; Bertelsmeier, 2021), records of species that were only transported to but not established within a particular region were aggregated with those of other species that established non-native populations within that region. Neglecting such differences in invasion capacities can bias perceptions of biosecurity threats as well as species’ viable geographic ranges, which are used widely in ecological niche modelling and climate-change predictions. Similar issues may arise from the aggregation of records from distinct climatic regions or biogeographic realms within coarse country-level boundaries (e.g., Australia, Brazil, China, and the USA in Bertelsmeier, 2021). A more detailed assessment— one addressing current gaps in information on species’ invasion capacities, barriers to invasion, and geographic scale—is therefore needed to clarify the global spread of alien ants as well as the biosecurity threats originating from and arriving at different parts of the world.

Here, we address key gaps in the knowledge of biological invasions by unintentionally introduced species. We (i) consolidate the total diversity of alien ant species and map their global distribution at unprecedented geographic resolution; (ii) clarify species’ capacities to overcome different barriers to invasion at regional and global scales; (iii) contrast the diversity, geographic spread, and habitat use among alien ants with different invasion capacities, as well as between alien ant species and all ant species; and (iv) test whether species that are spreading around the world are being detected at customs interceptions, and if there is any ecological bias to their detectability. Our test of habitat use as a basic correlate of invasion capacity is premised on evidence that ant assemblages using different vertical habitat strata are distinct in composition and ecological interactions (Ryder Wilkie et al., 2010; Wong & Guénard, 2017). Alien species occupying different strata may also use different introduction pathways (e.g., arboreal species in imports of fruits and wood; litter-dwelling species in imports of substrates) and therefore not be similarly detected in customs interceptions. Investigating alien species’ habitat may thus help identify major gaps in biosecurity measures.

Steps to accomplish the above aims are briefly described as follows (see Methods for additional details). We assembled data on ant species’ occurrences within their native and non-native ranges globally using the Global Ant Biodiversity Informatics (GABI) database (Guénard et al., 2017). The resultant dataset included 146,917 records corresponding to 17,948 occurrences of alien ant species across 602 non-overlapping regions (all areas where ants occur on Earth). We then organized the data according to the framework of Blackburn et al. (2011), which clarifies different invasion barriers. First, we summarized the native occurrences of all alien ant species across the 602 regions globally. Second, we used locality information to distinguish all non-native occurrences of alien ant species by invasion extent. We distinguished occurrences of species in customs interceptions (i.e., species overcoming geographical barriers to transport); occurrences of species established in indoor settings such as buildings and greenhouses (i.e., species overcoming demographic barriers and some environmental barriers to establishment); and occurrences of species established in outdoor settings including semi-natural and natural habitats (i.e., species overcoming most barriers to establishment). We also distinguished occurrences of species known to impact the ecosystems they invade as these are of utmost priority to invasive species management. Third, we identified the vertical habitat strata (i.e., the arboreal, ground-surface, or litter-and-soil strata; after Lucky et al., 2013) occupied by each alien ant species. The procedures allowed for comparing the diversity, geographic spread, and habitat use of alien ants with different invasion capacities globally. We also compared these to the native ranges and habitats of 15,487 ant species and subspecies (i.e., 99% of all extant taxa, Bolton, 2021; henceforth “all ant species”) to identify significant deviations from null expectations.

## RESULTS AND DISCUSSION

We identified a total of 522 alien ant species, which have been recorded as native in 522 regions and non-native in 489 regions, out of a total of 602 regions globally (suggesting that 113 regions are currently devoid of any alien ants based on current knowledge). Our data—available in Supporting information^1^—doubles the most recent estimate of alien ant diversity and improves the geographic resolution for mapping their global distributions (cf. 252 species distributed across 195 country-level units in Bertelsmeier, 2021). We placed alien ant species into four groups (Box 1) based on their maximum advancement—at the global scale—along a gradient of invasion extent and biosecurity threat (Fig. 1).

**Figure 1.**
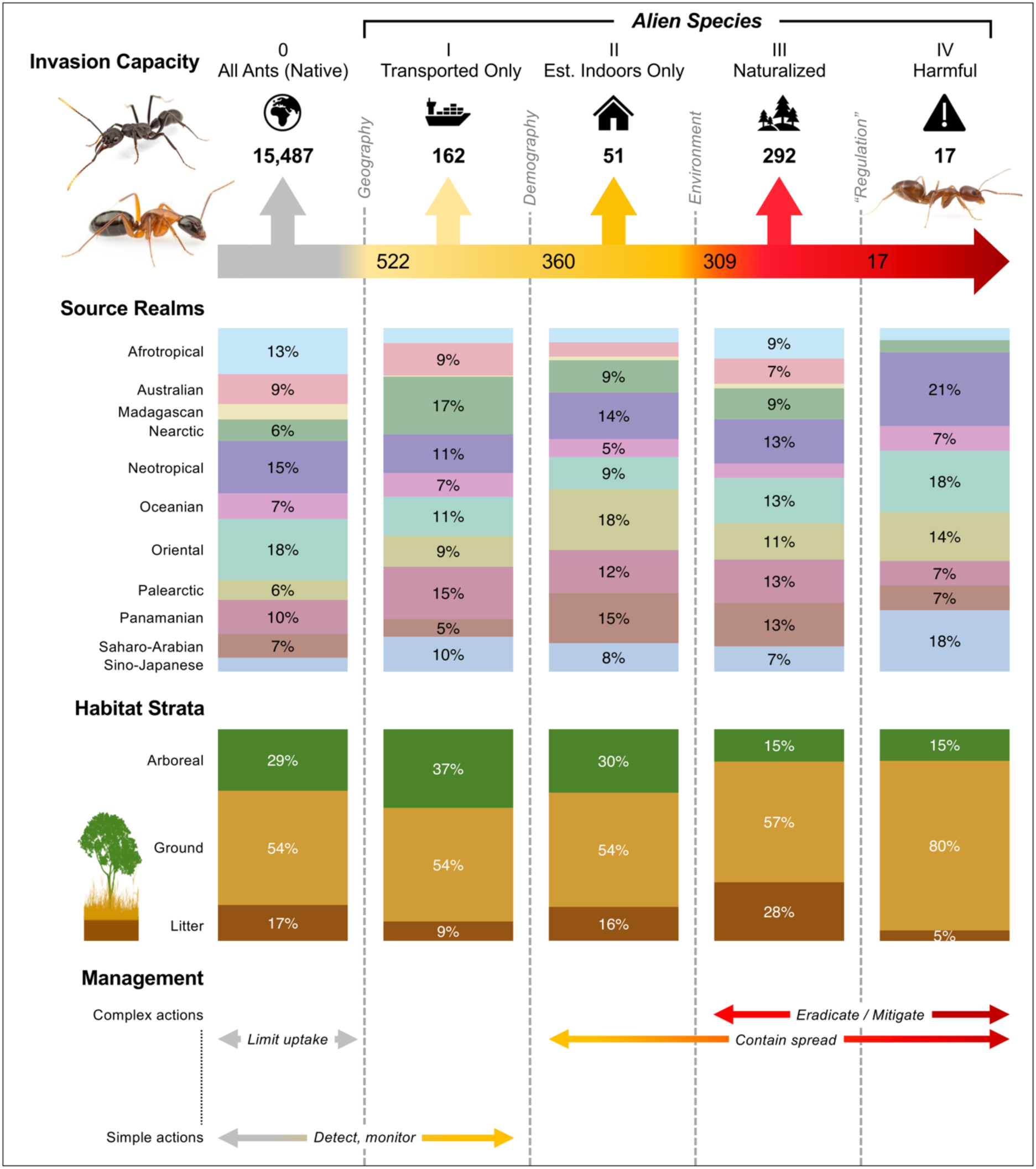
Differences among ant species’ invasion capacities, source realms, and vertical habitat strata globally. Top row (Invasion Capacity) summarizes the diversity of species in the global pool of ants [All Ants (Native)] as well as for 522 alien ant species in four levels of invasion capacity and biosecurity threat (Level I: Transported Only, Level II: Established Indoors Only, Level III: Naturalized, and Level IV: Harmful) that increase from left to right. Invasion capacities reflect species’ relative capacities to overcome successive barriers to invasion (vertical dotted lines) at the global scale. For species in each group, the second row (Source Realms) shows the distribution of their native regions across eleven major zoogeographic realms; the third row (Habitat Strata) shows their distribution across three major vertical habitat strata in terrestrial ecosystems; and the fourth row (Management) summarizes the most relevant actions for biosecurity management, which vary in complexity. Images by Alex Wild.

We found that the 522 alien ant species differed significantly (Chi-square tests, P<0.05) from the pool of all ant species globally in terms of their taxonomic composition, geographic origins, and use of vertical habitat strata (Fig. 1; Supporting information). This shows that alien ant species comprise a non-random subset of total ant diversity, with disproportionate contributions of species from distinct geographic source pools and vertical habitat strata. Moreover, we found that the taxonomic composition, geographic origins, and use of vertical habitat strata often varied significantly (Chi-square tests, P<0.05) among alien species with different invasion capacities at the global scale (Fig. 1; Supporting information). Different sets of factors may thus determine alien species’ capacities to overcome the geographical, demographic, and environmental barriers encountered during the successive invasion stages of transport, establishment, and spread (Blackburn et al., 2011). Alternatively, the patterns may also reflect differences in the likelihood of detecting different alien species in different settings and invasion stages (see “Blind spots in customs interceptions” below).

### Box 1. Invasion capacities of alien ants at global scale

Not all alien species possess equal capacities to invade. At the global scale, the invasion capacities of the 522 known alien ant species can be classified under four levels based on species’ maximum success in overcoming different barriers to invasion across 602 regions. The species in each level of invasion capacity are listed in Supporting information.

**Level I: Transported Only**

162 alien ant species have non-native records that only include individuals in customs interceptions at ports of entry. While such species can overcome barriers to their uptake and transport as stowaways on vehicles and cargo, successfully entering new destinations, they fail to overcome demographic and environmental barriers to their establishment of populations at these destinations. These ants therefore do not represent immediate threats to biosecurity, but their arrival should continue to be monitored.

**Level II: Established Indoors Only**

51 alien ant species can establish non-native populations within indoor settings only. Like Transported Only species, Established Indoors Only species can overcome barriers to their uptake and transport but upon arrival fail to overcome environmental barriers to establish non-native populations in the wild. These species can, however, overcome demographic barriers to establish (i.e., survive and reproduce) non-native populations in locations buffered from external environments, such as in buildings and greenhouses, where they may impact human activity as indoor pests. Given evidence that these alien species retain their demographic capacities under specific environmental conditions (i.e., indoor settings), they are ideal models for understanding environmental barriers to invasions.

**Levels III and IV: Naturalized and Harmful**

Of most concern to biosecurity, no less than 309 species—the majority of alien ant species globally—can establish non-native populations in outdoor settings in at least one region. Such ants possess substantially greater invasion capacities than the species described above. Not only can Naturalized and Harmful species overcome geographical, demographic, and environmental barriers to their establishment of non-native populations—even reaching high densities (Sanders et al., 2003), but doing so critically raises their chances for subsequent uptake and transport to expand their non-native ranges (Bertelsmeier & Ollier., 2021). Seventeen species (the Argentine Ant, Red Imported Fire Ant, African Big-Headed Ant, and others) are distinguished as Harmful (Level IV) based on evidence of their direct impacts on biodiversity and ecosystems, which often swiftly follow naturalization (Supporting information). However, this by no means implies that the remaining 292 Naturalized species (Level III) are of low concern; only that the impacts of these other species represent a critical knowledge gap, with some considered potential pests (e.g., the Ghost Ant, *Tapinoma melanocephalum*). It is unclear whether discrete ecological barriers regulate a transition from naturalized species to “harmful” or “invasive” species; the existence of a barrier also hinges on the precise definition and context of impact—a complex and disputed topic (Colautti & MacIsaac, 2004). Nonetheless, this barrier is tentatively termed “regulation” in Fig. 1.

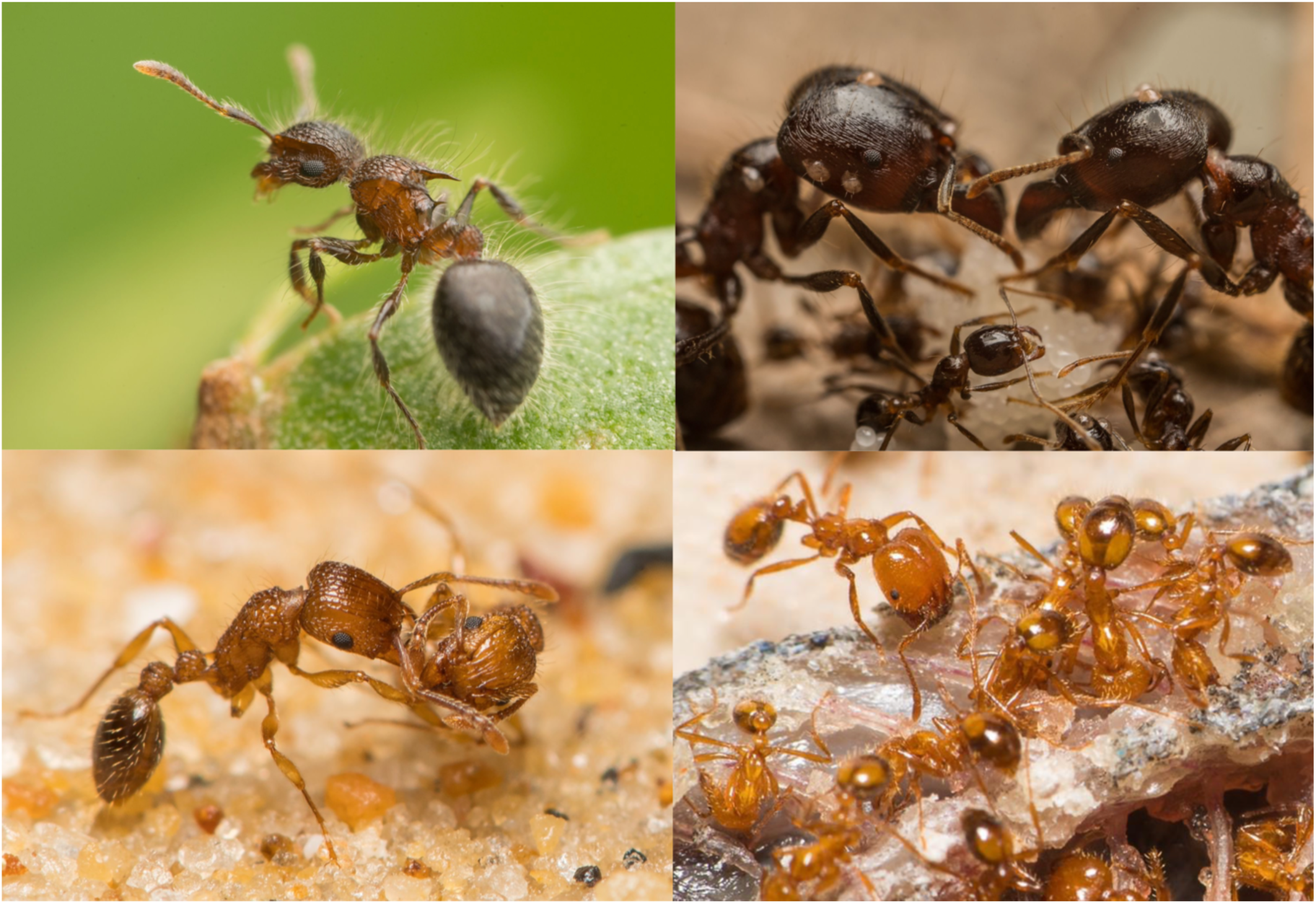

**Examples of alien ant species with different invasion capacities at the global scale**. Clockwise from top-left: *Meranoplus bicolor*, a Level I: Transported Only species recorded from customs interceptions in New Zealand and Louisiana; *Carebara diversa*, a Level II: Established Indoors Only species recorded from populations in a storage hangar in Italy; *Tetramorium bicarinatum*, a Level III: Naturalized species with non-native wild populations in multiple regions worldwide; *Solenopsis geminata*, a Level IV: Harmful species with non-native populations that have demonstrated harmful effects on native biodiversity in multiple regions worldwide. Images by Francois Brassard.

NB: The global classification of alien ants’ invasion capacities presented here is based on species’ *maximum* invasion extents across multiple regions globally. That is, the invasion capacities of individual species can vary at smaller scales, such as among different zoogeographic realms or individual regions. For instance, a species with an invasion capacity of “Level II: Established Indoors Only” globally may fail to establish indoors in or even enter some regions but not others; a species with an invasion capacity of “Level III: Naturalized” globally may fail to naturalize in or even enter some regions but not others. In addition to global-scale invasion capacities, realm- and region-scale invasion capacities were determined for alien species and used in analyses (Figs. 2,3;5).

**Figure 2.**
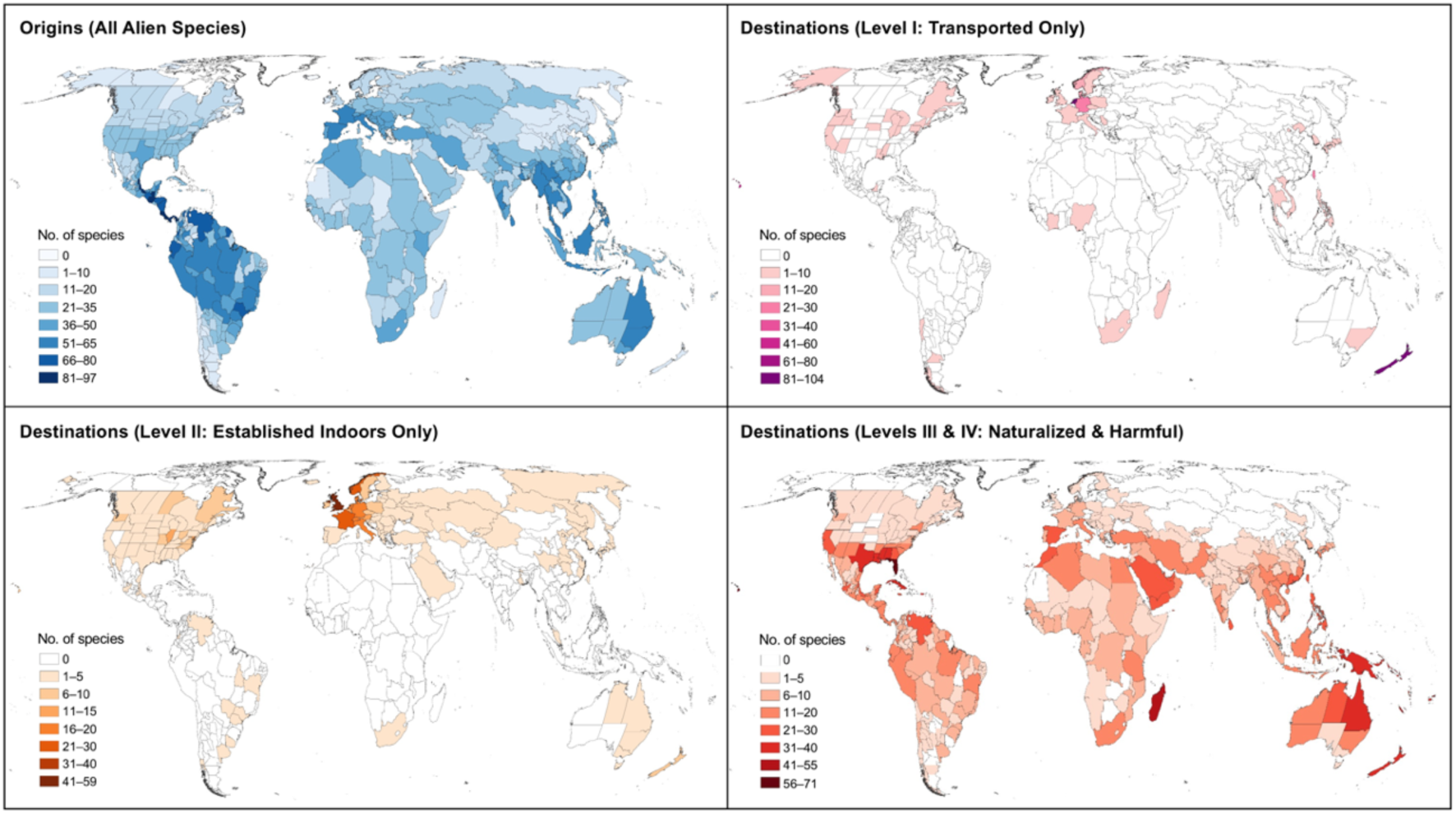
Global distributions (across 602 regions) of the origins of all alien ant species and the destinations of alien ant species of different invasion capacities (at region-scale). Region-scale invasion capacities are based on species’ maximum invasion extents within individual regions.

**Figure 3.**
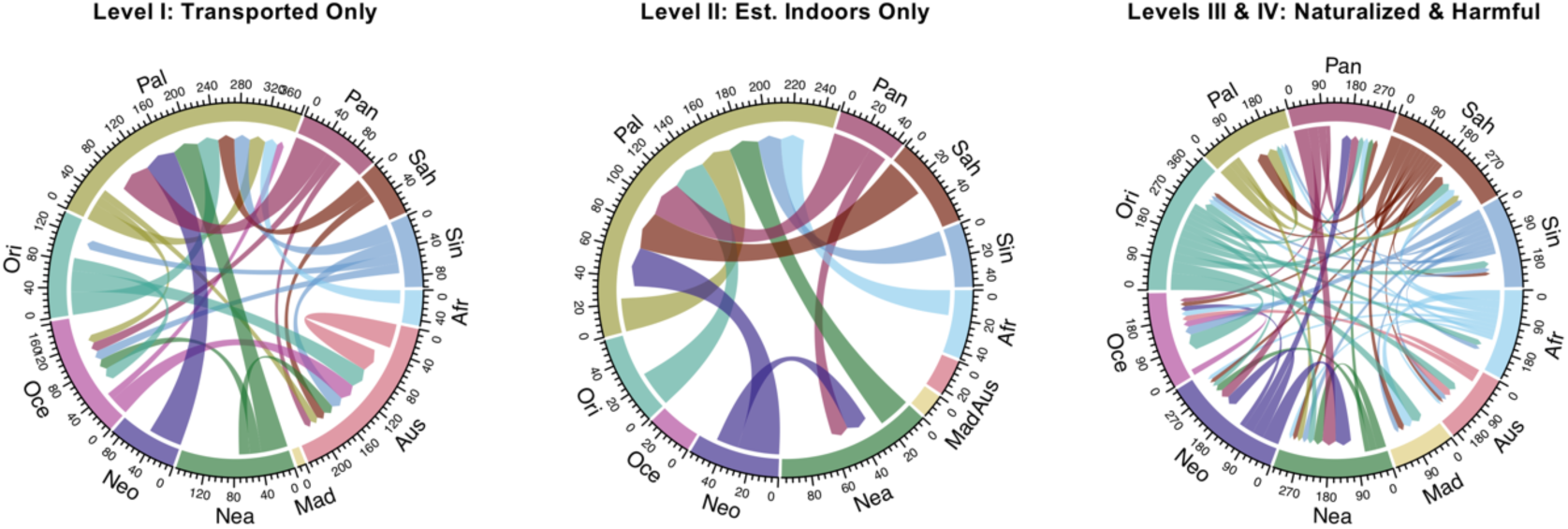
Flows in the diversity of alien ant species of different invasion capacities (at realm-scale) among eleven major zoogeographic realms. Realm-scale invasion capacities are based on species’ maximum invasion extents within individual zoogeographic realms. Flows with <10 species are not shown.

Global maps of alien species diversity and flows in the numbers of species exchanged between geographic realms are key information for understanding region-specific biosecurity risks and macroscale drivers of invasion (Van Kleunen et al., 2015). We found, however, that both the observed hotspots of alien ant species diversity and flows among geographic realms changed markedly when species of different invasion capacities were considered (Fig. 2–3).

### Invasion hotspots in individual regions and major zoogeographic realms

Across 602 potential destination regions globally, the hotspots where most alien ant species had established non-native populations within indoor settings were poorly correlated (*r*=-0.03) with those where most had overcome environmental barriers to establish non-native populations in the wild (Fig. 2). The highest diversity of naturalized and harmful alien ant species was found in the southern United States and various oceanic islands (Florida, Hawaii, and Mascarene Islands record 72, 58, and 55 species respectively). In contrast, regions in Europe showed the highest diversity of alien species with non-native populations that were restricted to indoor settings (United Kingdom, France, and Norway record 58, 26 and 23 species, respectively).

Distinctions in invasion capacities were likewise crucial when defining hotspots in alien ant diversity among Earth’s eleven major zoogeographic realms (after Holt et al., 2013). Although the Palearctic realm recorded the highest diversity of alien ant species overall (156 species), most records comprised species that were only transported to or which had established non-native populations only in indoor settings within this realm (Fig. 4). Fewer than half of all alien ant species in the Palearctic realm had overcome environmental barriers and naturalized—64 species—a number lower than that of substantially smaller areas such as the Saharo-Arabian and Madagascan realms (Fig. 4). In contrast, most alien ant species had naturalized in the Nearctic (108 species) and Oceanian (99 species) realms (Fig. 4).

**Figure 4.**
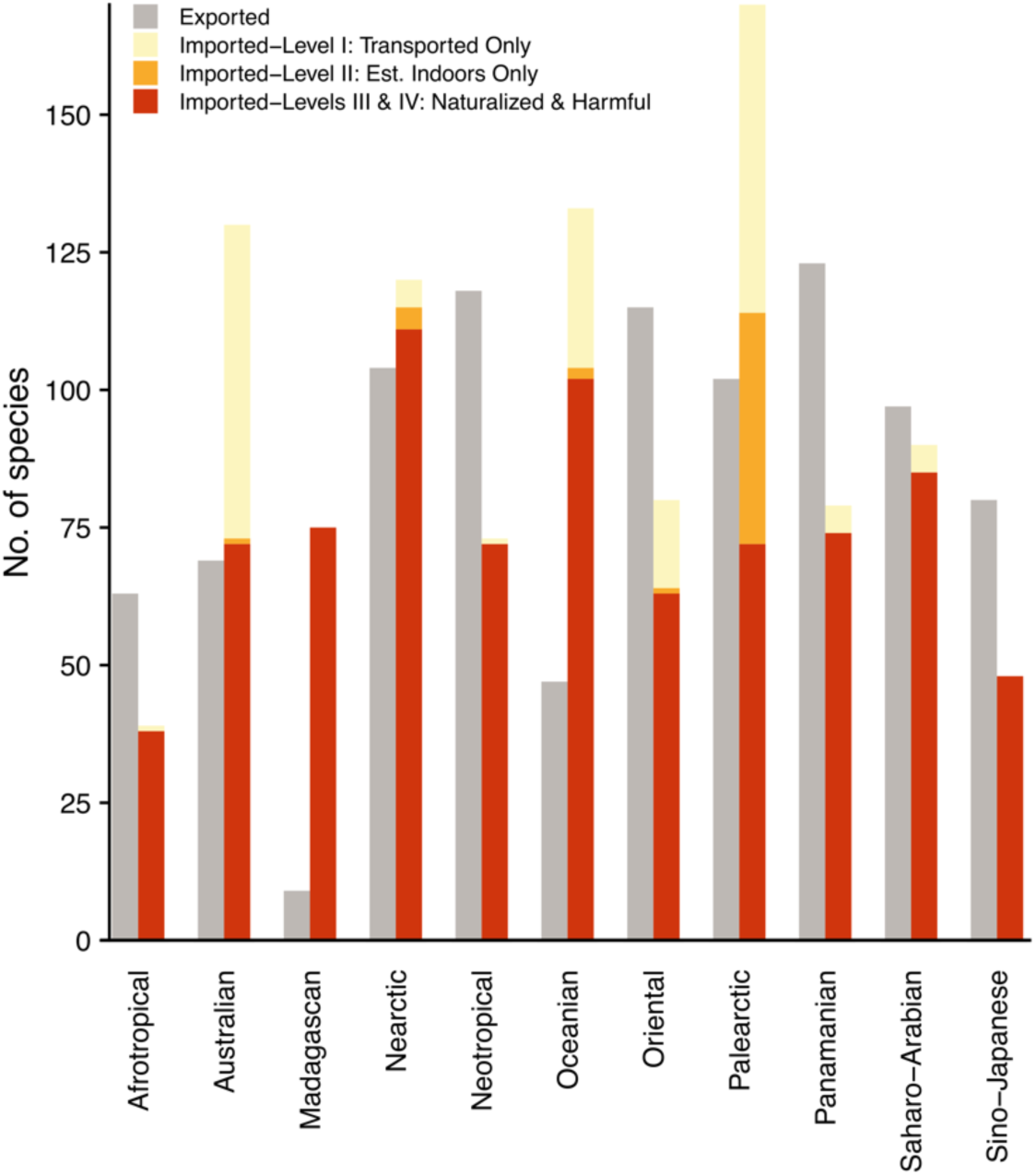
Numbers of alien ant species exported and imported by eleven major zoogeographic realms. Imported alien ant species are classified under different invasion capacities (at realm-scale), based on species’ maximum invasion extents within individual zoogeographic realms.

These global patterns align with those observed for naturalized plant and reptile species, which also peak in diversity in North America (Van Kleunen et al., 2015; Dawson et al., 2017). As postulated for many groups, an extensive history and high volumes of goods imported to the Nearctic from multiple regions globally could have substantially lowered geographical barriers to the transport of alien ant species to this realm (Hulme, 2009). Nonetheless, the large number of studies on biological invasions in the Nearctic (Pyšek et al., 2008) may also explain the many naturalized species detected. The high diversity of naturalized alien ant species in the Oceanian realm, which comprises numerous island systems, aligns with broad patterns observed in many groups (Moser et al., 2018), and may result from various ecological mechanisms that weaken barriers to the establishment of alien populations in islands, such as reduced biotic resistance and enemy release (Elton, 1958; Moser et al., 2018).

The relatively low diversity of naturalized alien ant species in the Palearctic realm is striking, given that geographical barriers to the transport of alien species to this realm would presumably have been extremely low for an extensive period throughout historic European colonization and maritime trade (Bertelsmeier et al., 2017). It also contrasts the high diversity of naturalized plant species reported in Europe (Van Kleunen et al., 2015). Environmental barriers to establishment, such as climatic or biotic resistance associated with the pervasive temperate climate across the Palearctic realm, may strongly mitigate ant invasions (Lach, 2021).

Two lines of evidence support the hypothesis that the Palearctic realm is distinctly characterized by a combination of low geographical barriers to transport but high environmental barriers to establishment. First, it supports the world’s highest diversity of alien ant species that establish non-native populations only in indoor settings (Fig. 4). Second, in comparison with the Palearctic realm, there is a disproportionately high diversity of species that have successfully naturalized in subtropical regions in the Nearctic realm such as Florida, Louisiana, and Mississippi (Fig. 4), which have likewise seen extensive historic maritime activity (Bertelsmeier et al., 2017), but which are subject to warmer climates. Alternatively, it is possible that some ant species thought to be native to the Palearctic realm may represent old introductions—in particular, from Mediterranean regions (Seifert et al., 2017)—that naturalized and spread very early throughout the realm. Additional information on population genetics would help discriminate such “cryptic” alien species.

### Flow patterns among major zoogeographic realms

Flow patterns of alien ant species exchanged between the eleven major zoogeographic realms changed substantially depending on the invasion capacities of the species considered (Fig. 3). Given their potential threats to biosecurity, we tested whether flows of naturalized ant species (including harmful ones) between individual realms were significantly above or below those expected from random (Fig. 5). We identified the Oriental realm as a significant exporter of naturalized and harmful ant species to six of ten other realms globally, exporting no less than 19 and as many as 52 species to these realms. (Fig. 5). Similarly, Turbelin et al. (2017) identified the Asia-Pacific region as the largest exporter of invasive alien species across multiple plant and animal groups, with most invasive alien species being native to China and India. While such patterns are attributed to complex and interacting biogeographic and economic factors, the patterns for ants nonetheless highlight a need for emphasized biosecurity measures to mitigate the spread of unintentionally introduced alien species from Asia.

**Figure 5.**
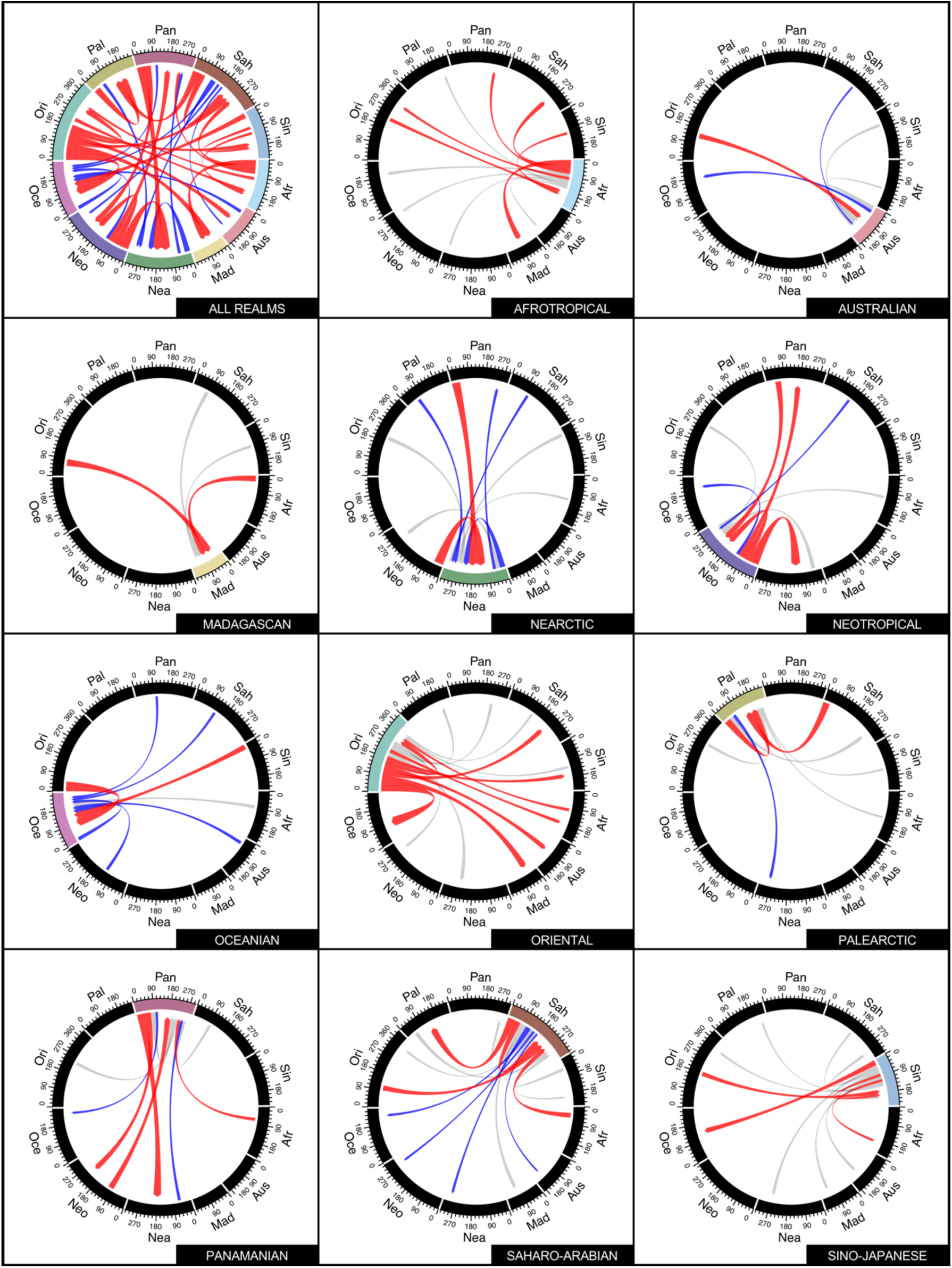
Flows in the diversity of alien ant species with an invasion capacity (at realm-scale) of Level III: Naturalized or higher exchanged among eleven major zoogeographic realms. First diagram shows flows for all realms; remaining diagrams show flows for individual realms. Diagrams illustrate results of resampling tests comparing the observed flows to 999 randomly drawn samples, where flows with values significantly higher than those drawn under random expectations (i.e., in the upper 2.5 percentile) are shown in red, and flows with values significantly lower than those drawn under random expectations (i.e., in the lower 2.5 percentile) are shown in blue; in diagrams for individual realms, flows with values which do not differ significantly from those drawn under random expectations are also shown in grey. Flows with <10 species are not shown.

### Blind spots in customs interceptions

The early detection and interception of alien species largely determines the success of biosecurity measures overall (Lodge et al., 2006). Our data showed that 291 alien ant species were recorded in customs interceptions globally (Supporting information). Remarkably, absent from this list were as many as two-thirds of all naturalized (198 species) species and three harmful species at the global scale (Supporting information). Many of these species likely evaded detection in repeated instances; as for numerous alien organisms, naturalized ant populations are unlikely to establish unless multiple colonization events have already occurred (Bertelsmeier & Ollier, 2021). While the general shortage of resources for screening at ports of entry surely accounts for this key blind spot in invasion management, biases in detection may also play a role.

Ecological studies show that systematic biases in detection can produce misestimates in the occurrences of species with (in)conspicuous traits or residing in (un)commonly sampled habitats (Gu & Swihart, 2004). Such biases in the detection of alien organisms are a less explored but nonetheless crucial aspect of biosecurity. We found a possible detection bias in customs interceptions toward arboreal ant species, which predominantly nest and forage in vegetation. Among the 291 alien ant species recorded in customs interceptions globally, the proportion of arboreal species was double and the proportion of litter-dwelling species less than half that among the 309 species capable of naturalizing (including harmful ones) at the global scale (Chi-square test, P<0.001; Supporting information). Furthermore, as many as 87% of the 198 species with naturalization capacity at global scale that were never recorded in customs interceptions were ground- and litter-dwelling (Supporting information). A bias toward arboreal species could occur if inspections of imports focused mainly on plant materials such as fruits and wood, as is often the case when screening for agricultural and domestic pests such as leaf-chewing or wood-boring insects (Kenis et al., 2007). Such measures place less emphasis on imported substrates and may therefore overlook myriad soil fauna that can disrupt native ecosystems (Ferlian et al., 2018).

Aside from potential detection biases, the high proportions of arboreal species among alien ants that are transported only (30%) as compared to those that are naturalized or harmful species (≤16%) (Fig. 1) may suggest that arboreal species have low success in establishing alien populations owing to environmental barriers. Arboreal alien ants may face direct biotic resistance from interspecific competition, which is often especially intense in vegetation, owing to the patchy distribution of limiting resources such as nest space and food (Mottl et al., 2021). Arboreal alien ants may also depend on mutualistic or facilitative interactions with plants or herbivorous insects that fail to establish in new regions owing to climatic factors.

### Conclusion

The spread of alien species globally is an ongoing ecological disaster. However, as shown here in the case of ants, not all alien species possess equal capacities to overcome geographical and ecological barriers and achieve successful invasions. Alien species with different invasion capacities originate from and accumulate at different regions globally, due to the influence of environmental barriers. Given the limited resources available for invasion management, recipient regions should prioritize biosecurity measures towards corresponding donor regions of species with high invasion capacities. Because the invasion capacity of species is driven in part by ecological attributes, such as affinity for different vertical habitat strata—biosecurity measures that do not take these differences into account (e.g., customs inspections) may achieve limited efficacy. Indeed, the fact that the majority of alien ant species being spread by humans have not been recorded by customs interceptions indicates that existing screening methods are woefully inadequate and need to be modified and/or supplemented by new approaches (e.g., Yasashimoto et al., 2021).

## METHODS

### Data compilation and organization

Data were compiled as part of the Global Ant Biodiversity Informatics (GABI) project; the details of data compilation are described fully in Guénard et al. (2017), and an update is presented here. In total, nearly two million records for nominal species were compiled from 10,342 scientific publications, as well as 82 and 16 public and private databases, respectively. Dubious and erroneous records were identified based on their mentions in the literature, cross-referencing when updates to checklists of species were available, and through direct communication with numerous experts since 2012.

Native and non-native ranges were identified for each species based on their mentions in the literature and the number of records present for the species (or in a few cases genera) within a particular biogeographic realm or part of it. Information about the locality and habitat of collection were also used to determine the native or non-native status of a species within a particular region. For instance, species encountered only within highly disturbed habitats within a particular region, one of the characteristics of tramp species (Passera, 1994), and with an uncertain native range were considered non-native within the region. For several species that displayed extensive and continuous distributions, some uncertainty was inevitable when demarcating the precise limits of their native and non-native ranges at the scale of individual regions; nonetheless these uncertainties would have limited influence on patterns of species distributions at the larger scale of zoogeographic realms. The resultant dataset, which included all ant species that had at least one non-native record globally, consisted of 146,917 records (including native and non-native records); 4,127 of these were subsequently identified as dubious records. We further summarized the data by geographic regions (Guénard et al., 2017) to yield 17,948 occurrences of alien ant species across 602 non-overlapping regions (i.e., all areas where ants occur on Earth).

Details about the collection locality, sampling method, or any other relevant information associated with each non-native record of a species were used to identify the geographical, demographic, and environmental barriers to invasion that the species had overcome (or failed to overcome) and therefore its invasion extent within that particular region (as described in Introduction). We compared the invasion extents of all non-native records for individual species to determine their invasion capacities (Level I: Transported Only, Level II: Established Indoors Only, or Level III: Naturalized) at the scale of individual regions and zoogeographic realms, as well as at the global scale. In the common case where a species had multiple non-native records with differing invasion extents at a given scale, we used a hierarchical approach and inferred the species’ invasion capacity from its maximum invasion extent at that scale.

To provide additional information to guide biosecurity measures, we scrutinized the pool of species with an invasion capacity of “Level III: Naturalized” and from these distinguished 17 “harmful” species for which there was evidence of their impacts on native biota in any of their invaded regions globally (Supporting information). We assigned the 17 species an invasion capacity of “Level IV: Harmful” in all non-native regions where they were previously recorded as “Level III: Naturalized.” We used this conservative approach because in the interests of biosecurity it would be safer to assume that a naturalized ant species could impact native biota if it had demonstrable impacts in other regions than to assume otherwise. For instance, naturalized ant species have shown to drive less obvious but nonetheless serious losses in functional diversity before more conspicuous declines in species numbers are observed (Wong et al., 2019).

It should be noted that while some studies of biological invasions have considered species to be “invasive” or “harmful” solely based on their geographic spread (Colautti & MacIsaac, 2004), such parameters are far more difficult to establish for ants because most parts of the world have received very limited sampling efforts (Guénard et al., 2012); even in frequently sampled areas, important habitat strata are often overlooked (e.g., Wong & Guénard 2017). Furthermore, species of some genera are extremely hard to identify in the absence of taxonomic expertise (e.g., *Cardiocondyla, Nylanderia, Pheidole*). Thus, we emphasized alien ant species’ impacts on native biota (as compared to their geographic spread) when flagging species that would be of the greatest priority for biosecurity. In this regard, our list of 17 harmful alien ant species differs slightly from the species listed in IUCN’s Global Invasive Species Database (GISD) (IUCN, 2021), which includes 19 species but for which, to the best of our knowledge, seven are lacking in evidence of impacts on native biota but may act as pests that are restricted to indoor environments or highly modified systems (see Supporting information). In addition, the GISD lacks some species, such as *Nylanderia fulva*, for which impacts on native biota have already been demonstrated (LeBrun et al., 2013).

We identified the vertical habitat strata used by each alien ant species based on a literature search as well as the genus-level classification of vertical habitat strata-use as reported in Lucky et al. (2013) with necessary updates to adhere to recent taxonomic changes. Three habitat strata were considered: the arboreal (“arboreal”), ground-surface (“ground”), and the litter-and-soil (“litter”) strata (Lucky et al., 2013). Ant species’ affinity for each of these strata was determined based on their foraging and nesting behaviors (see Lucky et al., 2013 for details). In coding the habitat strata of alien ant species in the present study, any species belonging to a genus that exclusively used a single stratum as reported in Lucky et al. (2013) was coded as that stratum; for instance, all species of *Hypoponera*, which nest and forage in leaf litter or soil, were coded as “litter.” For all other species, a literature search—typically using the web resources AntWiki (AntWiki, 2021) and AntWeb (AntWeb, 2021) to identify primary literature—was conducted to determine the stratum or strata used. If we found clear evidence for a species’ use of more than one stratum, all relevant strata (i.e., up to a maximum of three) were coded for that species. For instance, the Ghost Ant, *Tapinoma melanocephalum*, a species displaying extremely high environmental plasticity, was coded as “arboreal” and “ground” and “litter” as it has been observed foraging and nesting in all three strata (e.g., Sharaf et al., 2017). The vertical habitat strata of alien ant species in our study are provided in Supporting information.

### Data analysis

We used Chi-square tests to compare the frequency distributions of the native zoogeographic realms and vertical habitat strata between alien ant species in the four different invasion capacities (defined at the global scale). We also compared each of these to the frequency distributions of the native zoogeographic realms of all ant species (using data from GABI, Guénard et al., 2017) and the vertical habitat strata of all ant genera (using updated data from Lucky et al., 2013). We characterized flows of naturalized ant species (including harmful ones) within and between different zoogeographic realms, and used resampling tests (after Van Kleunen et al., 2020) to assess whether the observed flows were statistically larger or smaller than expected. The resampling tests involved comparing the observed flows to flows based on 9,999 random draws from the full list of naturalized ant species. If the observed number was in the upper or lower 2.5% quantiles of the resampled values, the flows were considered to be significantly higher or lower, respectively, than expected.

## ACKNOWLEDGEMENTS

We thank Alex Wild and Francois Brassard for providing photographs of ant species, and Mark Van Kleunen for sharing scripts for the resampling analyses. This work was supported by the Japan Ministry of the Environment (Environment Research and Technology Development Fund no. 4-1904 to E.P.E.).

## Footnotes

1 Supporting information will be made available upon publication and is not included in this preprint.

